# Brain strain rate response: addressing computational ambiguity and experimental data for model validation

**DOI:** 10.1101/2022.02.04.478773

**Authors:** Zhou Zhou, Xiaogai Li, Yuzhe Liu, Warren N. Hardy, Svein Kleiven

## Abstract

Traumatic brain injury (TBI) is an alarming global public health issue with high morbidity and mortality rates. Although the causal link between external insults and consequent brain injury remains largely elusive, both strain and strain rate are generally recognized as crucial factors for TBI onsets. With respect to the flourishment of strain-based investigation, ambiguity and inconsistency are noted in the scheme for strain rate calculation within the TBI research community. Furthermore, there is no experimental data that can be used to validate the strain rate responses of finite element (FE) models of the human brain. Thus, the current work presented a theoretical clarification of two commonly used strain rate computational schemes: the strain rate was either calculated as the time derivative of strain or derived from the rate of deformation tensor. To further substantiate the theoretical disparity, these two schemes were respectively implemented to estimate the strain rate responses from a previous-published cadaveric experiment and an FE head model secondary to a concussive impact. The results clearly showed scheme-dependent responses, both in the experimentally determined principal strain rate and FE model-derived principal and tract-oriented strain rates. The results highlight that cross-scheme comparison of strain rate responses is inappropriate, and the utilized strain rate computational scheme needs to be reported in future studies. The newly calculated experimental strain rate curves in the supplementary material can be used for strain rate validation of FE head models.

## 1 Introduction

Traumatic brain injury (TBI), defined as an alteration in brain function or other evidence of brain pathology caused by an external force [1], is a major global public health threat. In the United States, around 61000 people died from TBI-induced injuries in 2019 [2], and in the European Union, there were approximately 82000 TBI-associated deaths in 2012 [3]. Despite collective efforts from the scientific community, clinical services, and policymakers to prevent the occurrence and mitigate the detriment of TBI, no clear improvement has been noted over the past two decades [4]. Efforts to better decipher brain injury pathogenesis and develop more effective head protective strategies are vital to resolve this concerning epidemic.

Although the mechanobiological pathway from external insult, localized tissue responses, and resultant brain injury remains to be better explored, it has been generally understood that both strain and strain rate are crucial factors for TBI onsets [5]. For example, through relating controlled mechanical inputs to neuromorphological changes in a three-dimensional (3D) *in vitro* compression model, Bar-Kochba et al. [6] found that the localization of neurite blebbing correlated with shear strain along the axial direction of neurons, while neurite swelling was highly loading-rate dependent. Montanino et al. [7] utilized a composite finite element (FE) model of a single axon to predict the responses of axonal subcomponents secondary to tensile elongation, revealing a dependency of axolemma deformation on both the magnitude and the rate of imposed loadings. Hajiaghamemar and Margulies [8] found FE model-derived strain and strain rate along the fiber tracts spatially aligned with the pattern of histology-identified axonal injury in the pig brains after rapid head rotations. These experimental and computational findings collectively underscored the importance of monitoring both the strain and strain rate during head impacts.

Various strain-based metrics have been proposed for TBI analyses, such as maximum principal strain (MPS) [9-16], cumulative strain damage measure (CSDM) quantifying the volume fraction of brain experiencing strain levels over a pre-defined threshold [17, 18], maximum tract-oriented strain (MTOS) measuring the stretch along axonal fiber tract [19-21]. Strain-based measurements have been experimentally quantified by Hardy et al. [22] (recently reanalysed by Zhou et al. [23]) based on MPS (0.07-0.22) in cadaveric experiments with loading conditions close to traumatic levels, and by Knutsen et al. [24] based on MPS (0.02-0.05) and MTOS (0.005-0.03) in voluntary loading scenarios. These strain-based metrics were also used to establish kinematic-based brain injury risk functions with the aid of FE head models. For example, the Brain Injury Criteria (BrIC) [25] was proposed based on the dependency of MPS and CSDM on the angular velocity observed from two FE models of human brain [17, 26]. Discussions are ongoing to potentially adopt this newly proposed BrIC into the New Car Assessment Program in the vehicle safety standards [27].

Compared with the large body of strain-based studies, there are relatively fewer investigations on brain strain rate. It is ubiquitous in the literature that the strain rate values are directly reported without any clarification on how they are calculated [9, 28-33]. Even when limiting to the handful of studies with the strain rate computational schemes elaborated, alarming inconsistency existed. As typical practices, the strain rate is either cacluated from the rate of deformation tensor [34, 35] or computed as the time derivative of strain [36, 37]. However, little knowledge exists on how the computational schemes affect the resultant strain rate values.

Today, there is no existing experimental brain strain rate dataset with loading conditions close to traumatic levels and hence the accuracy of the FE model-derived strain rate response remains unknow. Commonly, the FE model of human brain is validated against the brain-skull relative motion and/or brain strain and then directly used for strain rate prediction. As highlighted by Zhou et al. [38] and Zhao and Ji [39], if the model is evaluated against an experimental parameter that is not the parameter of interest itself, it is suspected that numerical error might be introduced and propagated through the chain of calculation from the evaluated parameter to the parameter of interest (e.g., the relationship between brain motion/strain and brain strain rate). For an FE model with intended usage of strain rate prediction, a direct evaluation of brain strain rate response is thus preferred.

The current study endeavoured to address the ambiguity relevant to the calculation of brain strain rate and provide experimental data for strain rate evaluation of FE models of the human brain. Thus, a theoretical clarification of two commonly used strain rate computational schemes was presented, in which the strain rates were computed as the time derivatives of strains of interest (referred to as scheme 1), and components of interest of the rate of deformation tensor (referred to as scheme 2), respectively. These two schemes were then implemented to calculate the experimental principal strain rate and computational principal and tract-oriented strain rates, through which the distinction between these two schemes was substantiated. The newly calculated experimental strain rate curves can be served as a reference to evaluate the strain rate responses of FE models.

## 2 Methods

### 2.1 Theoretical clarification

This section presents a continuum mechanical description of the two strain rate computational schemes (i.e., scheme 1 and scheme 2), in which the calculation of two specific strain rates are elaborated (i.e., the first principal strain rate and the tract-oriented strain rate). Two strain rates in combination with two computation schemes thus yields 4 types of strain rates, all of which are used as response variables in TBI studies. The mathematical symbols used throughout the text are clarified below: 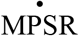 and 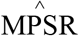 represent the maximum first principal strain rates computed by scheme 1 and scheme 2, respectively; 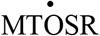 and 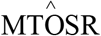 namely denote the maximum tract-oriented strain rates following scheme 1 and scheme 2.

As is the common context of these two schemes, the motion **φ**(**X**, *t*) maps the position vector of every point in the brain in the undeformed configuration (**X**) (i.e., **X** ={*X*_1_ *X*_2_ *X*_3_}) to its deformed configuration (**x**) at time t (i.e., **x**(*t*) = {*x*_1_(*t*) *x*_2_ (*t*) *x*_3_(*t*)}). The temporal derivative of **φ**(**X**, *t*) at fixed position **X** represents the velocity, writing as **v**(**X**, *t*) = {*v*_1_(*t*) *v*_2_ (*t*) *v*_3_(*t*)}.

#### 2.1.1 Scheme 1: Time derivative of strain

Scheme 1 calculates the strain rates as the time derivatives of the first eigenvalue or the component of interest of the strain tensor **E**(*t*). Given that **φ**(**X**, *t*) consists of both deformation and rigid-body motion, the deformation gradient **F**(*t*) is introduced, isolating the deformation through mapping line elements (d**X**) in the reference configuration into its counterpart (d**x**) in the current configuration.

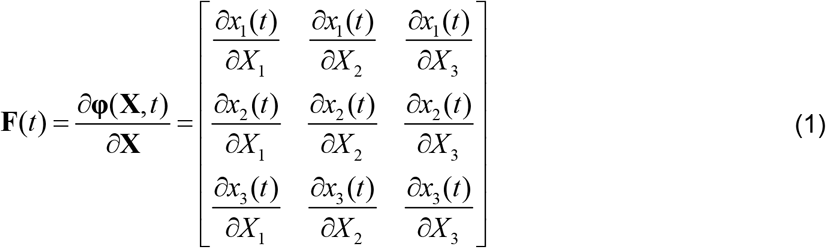

The Green-Lagrange strain tensor is then calculated as

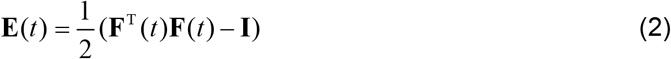

where the superscript T represents the transpose operation and **I** is the identify tensor.

The first principal strain (PS(*t*)) is computed as the maximum eigenvalue of **E**(*t*) and the tract-oriented strain (TOS(*t*)) is obtained by projecting **E**(*t*) along the real-time fiber orientation (**a**(*t*)) (equation 3) [40]:

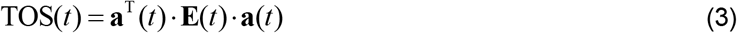

Resultantly, the first principal strain rate 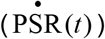 and tract-oriented strain rate 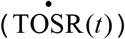 were computed as the derivative of PS(*t*) and TOS(*t*) with respect to time (equations 4 and 5), respectively. Of note, a five-point stencil neural network derivative function was used for all the derivative operations, same as the approach used in previous studies [41-43].

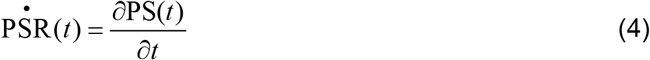

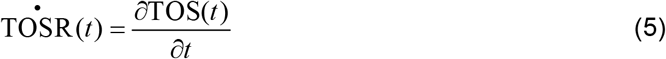

The maximum value of 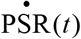 and 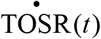 across the simulated time domain was identified (i.e., 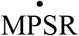 and 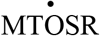, respectively). Scheme 1 has been used to calculate the maximum principal strain rate in the brain [8, 19, 36, 41-49] and maximum tract-oriented strain rate in the white matter (WM) [8, 19, 41, 42, 44-46, 48].

#### 2.1.2 Scheme 2: Rate of deformation tensor

Scheme 2 calculates strain rate as the first eigenvalue or the component of interest of the rate of deformation tensor **d**(*t*). Of note, the rate of deformation tensor is synonymous with rate of strain tensor [50]. In contrast with the deformation gradient **F**(*t*) based on displacement field, velocity gradient **l**(*t*) is derived from velocity field **v**(**X**, *t*) by taking its spatial gradient according to equation 6:

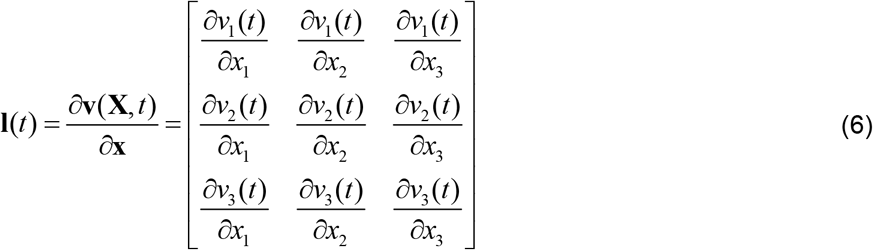

The symmetrical part of **l**(*t*) is defined as the rate of deformation tensor **d**(*t*) :

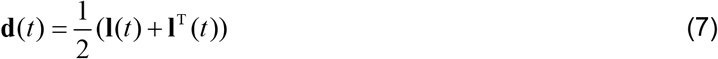

The first principal strain rate 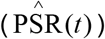 and tract-oriented strain rate 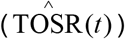 were respectively computed as the maximum eigenvalue of **d**(*t*) and the projection of **d**(*t*) along the real-time fiber orientation (**a**(*t*)) (equation 8).

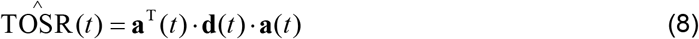

The time-accumulated peaks of 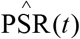 and 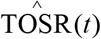 were determined and labelled as 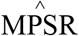 and 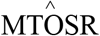, respectively. Scheme 2 has been previously used to extract the strain rate in the brain [35, 51] and other biological tissues (e.g., leg muscles [52-55], myocardium [56-58]).

While collectively considering these two schemes, it is worth mentioning that the material time derivative of the Green-Lagrange strain tensor (i.e., **E** in scheme 1) is the pull-back of the rate of deformation tensor (i.e., **d** in scheme 2) (equation 9) [50, 59]. 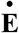 in equation 9 is termed as material strain rate tensor in the literature [50] and, to the best of the authors’ knowledge, has also been used to inform the computation of brain strain rate in two studies from one research group [60, 61]. Given its relatively limited usage within the TBI community, this scheme is thus not incorporated in the current study.

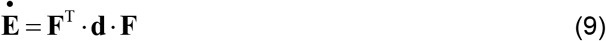

### 2.2 Experimental strain rate

To substantiate the disparity between scheme 1 and scheme 2 and provide experimental data for strain rate evaluation of FE brain model, we computed the principal strain rates in cadaveric brains using both schemes. In Hardy et al. [22], thirty-five impacts were conducted on eight inverted and perfused head-neck complexes of post-mortem human subjects, in which the motion of the radiopaque neutral density targets (NDTs) implanted within the cadaveric brain was measured using a high-speed, bi-planar X-ray system with a temporal resolution of 1 ms. To expedite the estimation of local responses, Hardy et al. [22] deployed 7 NDTs in a cluster configuration with the center target being 10 mm from the remaing six counterparts, roughly occupying 1.5 ml of tissue. As represented by C1 and C2 in Figure 1A, two NDT clusters were implanted within each cadaveric brain.

**Figure 1.**
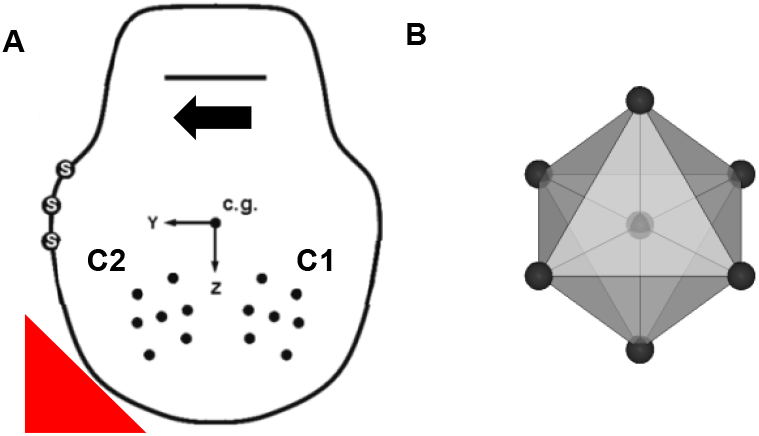
(A) Representative sketch of NDT cluster implanting schemes. For a given head, two NDT clusters were implanted as cluster 1 (C1) and cluster 2 (C2). (B) Strain-rate estimation with the NDTs as black spheres while tetrahedron elements in grey and mesh line in black. To visualize the center NDT, all tetrahedron elements are shown in translucency.

Following the selection criteria proposed by Zhou et al. [38], 15 NDT clusters from 14 impacts with both impact kinematics and motions of all 7 associated NDTs recorded over 40 ms were chosen to calculate the principal strain rates. As illustrated by a representative cluster in Figure 1B, FE nodes were generated based on the NDT starting positions. Based on the NDT deploying characteristics in Hardy et al. [22], 8 tetrahedron elements were developed for each cluster by connecting each NDT to its neighbouring counterparts [23]. The tracked motions were used to excite the tetrahedron model, which was solved by the LS-DYNA software (Livermore Software Technology Corporation, LSTC). From the completed simulation, the first principal strain rates were respectively extracted as the time derivatives of the first principal strain (i.e., 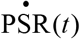 for the scheme 1) and maximum eigenvalue of the rate of deformation tensor (i.e., 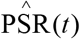 for scheme 2). The strain rates of all available tetrahedron elements were averaged to provide a general estimation of the strain rate within the volume occupied by the NDT cluster. The maximum values of 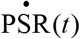 and 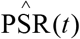 were identified (i.e., 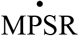 and 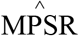). Due to the unavailability of fiber orientation information of these cadavers, it is infeasible to compute tract-oriented strain rate from this experimental data.

### 2.3 Computational strain rate

#### 2.3.1 The ADAPT model

We next implemented scheme 1 and scheme 2 to calculate the principal strain rate and tract-oriented strain rate from an anatomically detailed head model (i.e., the ADAPT model in Figure 2). The ADAPT model was developed previously at KTH Royal Institute of Technology in Stockholm using the LS-DYNA software [62]. The model includes the brain, skull, meninges (i.e., dura mater, pia mater, falx, and tentorium), subarachnoid cerebrospinal fluid (CSF), ventricle, and sinuses (i.e., superior sagittal sinus and transverse sinus). As is the structure of interest in the current study, the brain was further grouped with explicit representations of various components, including four lobes of the cerebral gray matter (GM) (Figure 2A), cerebellar GM, cerebral WM, cerebellar WM, and brainstem (Figure 2B). The element numbers of brain and WM are 1513198 and 213321, respectively. A second-order Ogden hyperelastic constitutive model combined with six-order Prony series was employed to describe the nonlinearity and viscoelasticity of brain material [63]. The biofidelity of the model has been evaluated against experimental data of intracranial pressure [64], brain-skull relative displacement [22], and brain strain [23]. The strain rate responses of the model were additionally evaluated by the newly calculated experimental strain rate data in Appendix.

**Figure 2.**
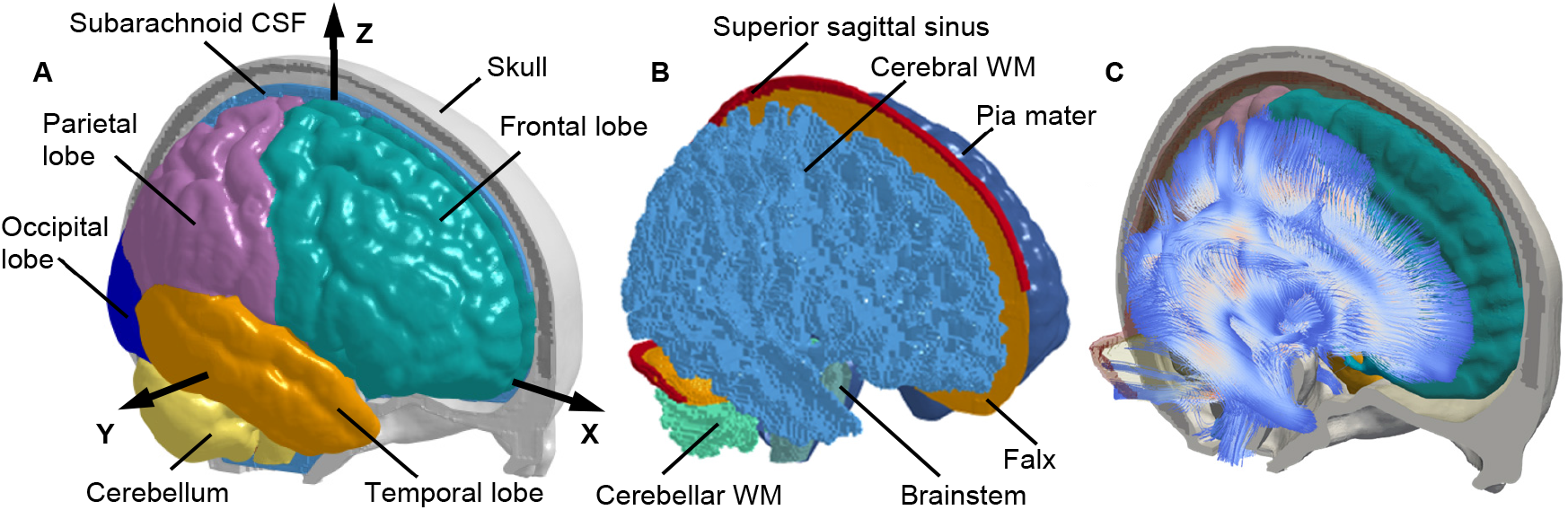
Finite element model of the human head with embedded truss elements representing te white mater fiber tracts. (A) Finite element model of the human head with the skull open to expose the subarachnoid cerebrospinal fluid (CSF) and brain. A skull-fixed coordinate system and corresponding axes are illustrated with the origin at the head’s centre of gravity. (B) Brain components of cerebral white matter (WM), cerebellar WM, brainstem, pia mater, falx, tentorium, superior sagittal sinus, and transverse sinus. (C) Brain with fiber tracts.

To extract tract-related responses, it is imperative to monitor the real-time direction of fiber tracts during the impact [40]. Thus, the primary orientation of the WM fiber tracts was extracted from the ICBM DTI-81 atlas [65] and then embedded within the WM elements in the form of truss elements [40]. Through this, potential variation in fiber orientation during the impacts was reflected by the temporal direction of the truss element, which was updated at each solution cycle of the time-marching simulation. These temporally dynamic fiber orientation informed the computation of tract-oriented strain rate.

#### 2.3.2 Impact simulation

The ADAPT model was employed to simulate a concussive impact, through which both the maximum principal strain rates and maximum tract-oriented strain rates were computed using scheme 1 and scheme 2. The impact loading was measured by instrumented mouthguard from a collegiate American football player with self-reported post-concussive symptoms [66]. Both translational and rotational acceleration profiles (Figure 3) were applied to a node located at the head’s centre of gravity and constrained to the rigidly modelled skull. The model responses were output at every 0.1 msec, the same as the output interval in [41].

**Figure 3.**
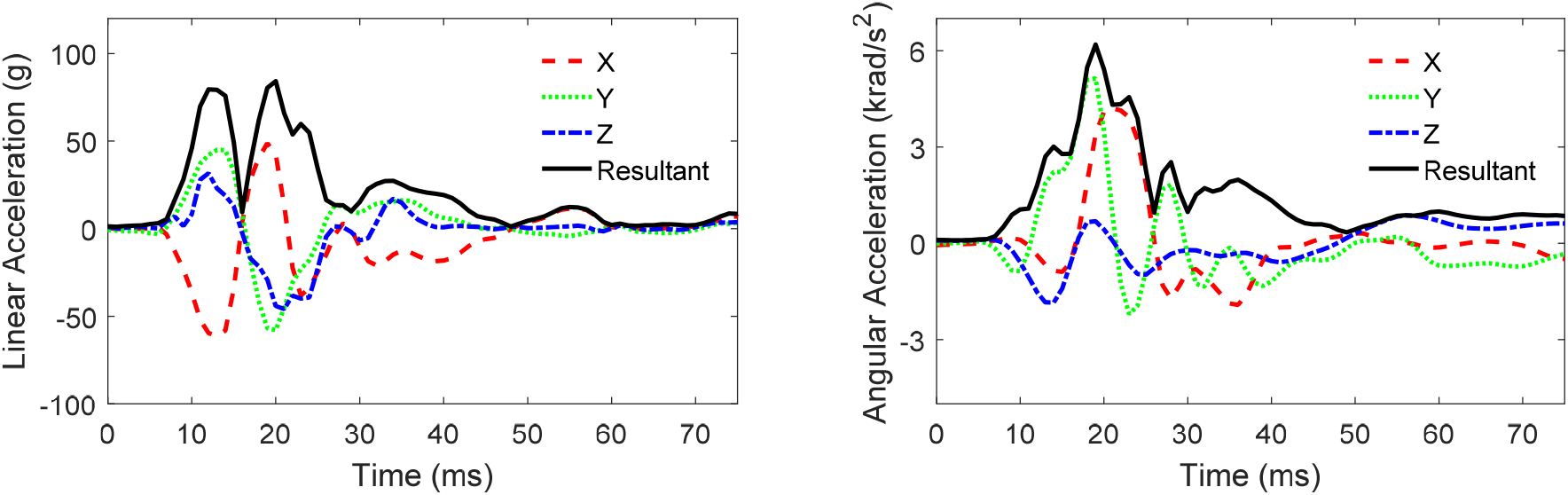
Head model loading conditions for one concussive impact with the kinematics measured from instrumented mouthguard. Note the X, Y, and Z axes are the same as those in the skull-fixed coordinate system in Figure 2A.

Once the simulation was complete, the strain tensor and rate of deformation tensor of all brain elements and the temporal orientation of the truss elements embedded within the WM elements were obtained. For each brain element, the first principal strain rates were calculated as the time derivatives of the first principal strain (i.e., scheme 1) and the first eigenvalue of the rate of deformation tensor (i.e., scheme 2) with their maximum values across the whole impact reported (i.e., 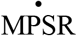 and 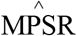). The tract-oriented strain rates were additionally computed for all WM elements. In scheme 1, the strain tensor of each WM element was iteratively resolved along the temporal direction of its embedded truss element (i.e., equation 3) with the outcome being discretely differentiated between time points to obtain the tract-oriented strain rate (i.e., equation 5). In scheme 2, the rate of deformation tensor of each WM element was projected along the real-time direction of its embedded truss element (i.e., equation 8). For both schemes, the maximum values of the tract-oriented strain rates across the whole impact were reported, i.e., 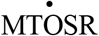 and 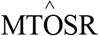.

#### 2.3.3 Data analysis of computational strain rate responses

To quantify the cross-scheme disparity, relative differences in strain rate peaks computed by scheme 1 and scheme 2 were computed for all brain elements based on the principal measures and for all WM elements based on the tract-oriented measures, respectively. To test whether cross-scheme strain rate results were readily translatable, linear regression analyses were conducted between 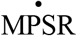 and 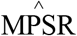, and between 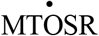 and 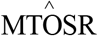, respectively. In addition to the correlation coefficient (r), the root-mean-square error normalized by the mean value of strain rate peaks calculated by scheme 1 (i.e., NRMSE) was reported. The threshold for significance was *p*<0.05.

## 3 Results

### 3.1 Experimental principal strain rates

Experimental principal strain rates computed by both schemes for all 15 clusters are depicted in Figure 4. Using scheme 1, the time history curves of the principal strain rate were generally biphasic with both positive and negative phases. When switching to scheme 2, the strain rate curves were monophasic with purely positive values. The strain rate responses of all 15 clusters attained the peak values within the pulse durations except for C241-T5 C2 and C393-T1 C1 solved by scheme 2. The strain rate peaks along with experimental kinematic peaks are summarized in Table 1. The maximum values ranged from 9.3 s^-1^ (C241-T6 C2) to 25.8 s^-1^ (C380-T4 C1) in scheme 1 and spanned from 15.6 s^-1^ (C380-T2 C1) to 35.6 s^-1^ (C064-T1 C1) in scheme 2. Note that the experimental principal strain rate-time history curves in Figure 4 are available in the supplentary materials.

**Table 1.**
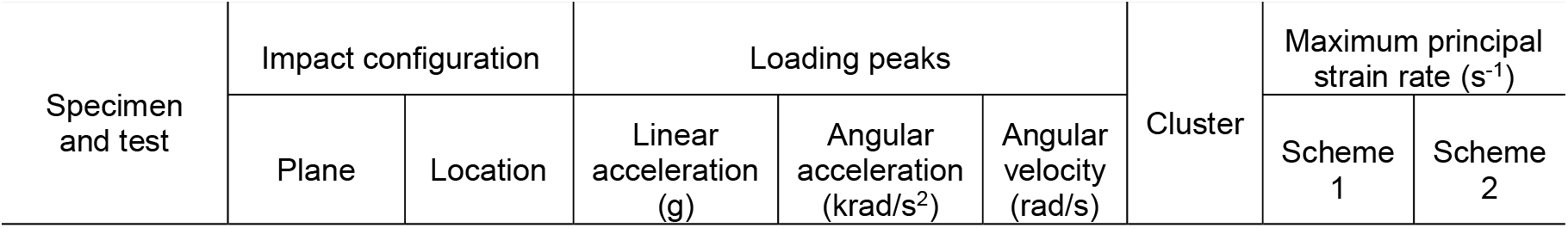
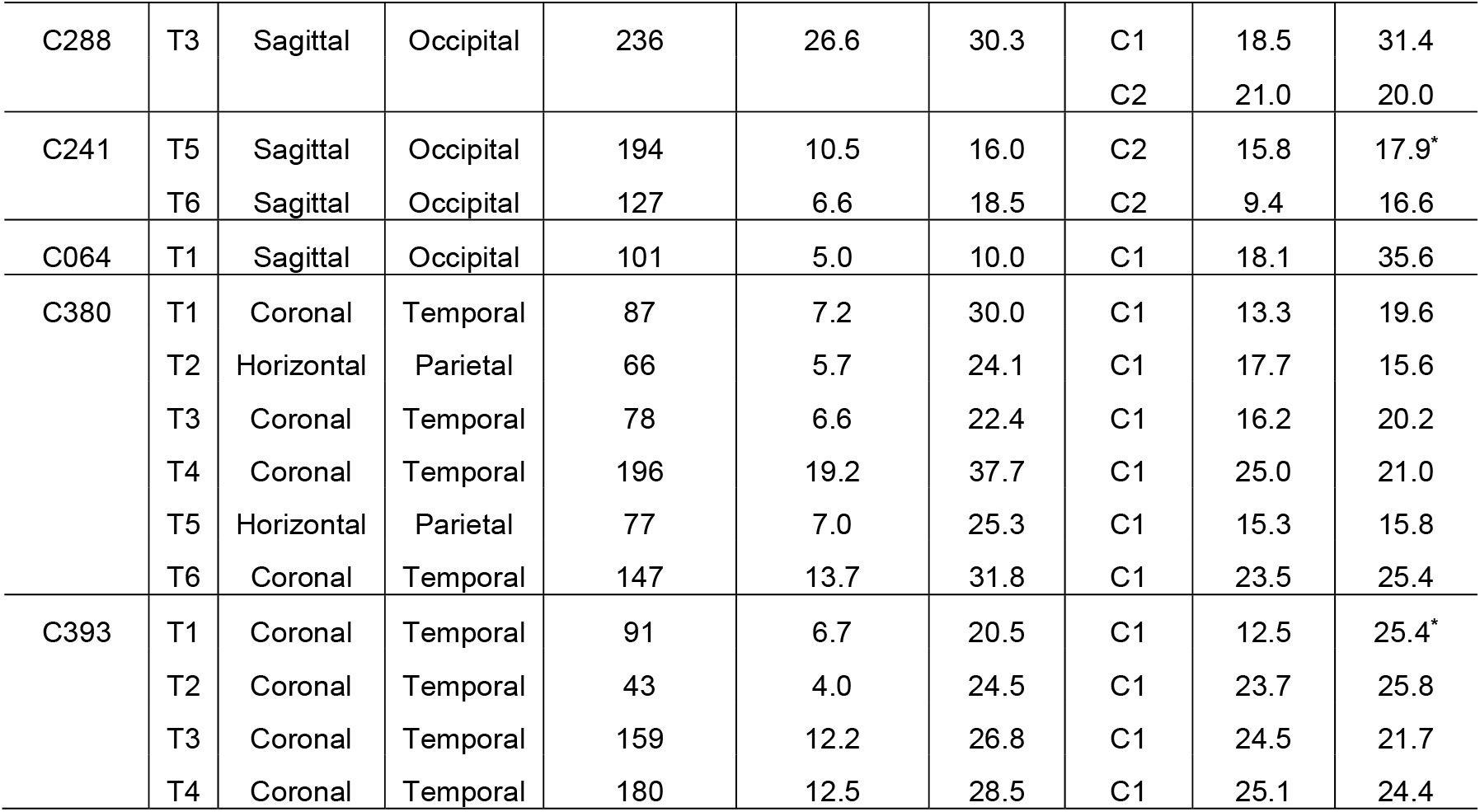
Summary of test configurations of 14 impacts and strain rate responses of 15 NDT clusters. Note that the loading peaks are calculated based on the resultant kinematics and the superscript **\*** indicates the strain rate does not reach the peak value within the available time duration.

| Specimen and test | Impact configuration | | Loading peaks | | | Cluster | Maximum principal strain rate ($s^{-1}$) | |
| --- | --- | --- | --- | --- | --- | --- | --- | --- |
|  | Plane | Location | Linear acceleration (g) | Angular acceleration (krad/s<sup>2</sup>) | Angular velocity (rad/s) |  | Scheme 1 | Scheme 2 |

|  |  |  |  |  |  |  |  |  |  |
| --- | --- | --- | --- | --- | --- | --- | --- | --- | --- |
| C288 | T3 | Sagittal | Occipital | 236 | 26.6 | 30.3 | C1 | 18.5 | 31.4 |
|  |  |  |  |  |  |  | C2 | 21.0 | 20.0 |
| C241 | T5 | Sagittal | Occipital | 194 | 10.5 | 16.0 | C2 | 15.8 | 17.9* |
|  | T6 | Sagittal | Occipital | 127 | 6.6 | 18.5 | C2 | 9.4 | 16.6 |
| C064 | T1 | Sagittal | Occipital | 101 | 5.0 | 10.0 | C1 | 18.1 | 35.6 |
| C380 | T1 | Coronal | Temporal | 87 | 7.2 | 30.0 | C1 | 13.3 | 19.6 |
|  | T2 | Horizontal | Parietal | 66 | 5.7 | 24.1 | C1 | 17.7 | 15.6 |
|  | T3 | Coronal | Temporal | 78 | 6.6 | 22.4 | C1 | 16.2 | 20.2 |
|  | T4 | Coronal | Temporal | 196 | 19.2 | 37.7 | C1 | 25.0 | 21.0 |
|  | T5 | Horizontal | Parietal | 77 | 7.0 | 25.3 | C1 | 15.3 | 15.8 |
|  | T6 | Coronal | Temporal | 147 | 13.7 | 31.8 | C1 | 23.5 | 25.4 |
| C393 | T1 | Coronal | Temporal | 91 | 6.7 | 20.5 | C1 | 12.5 | 25.4* |
|  | T2 | Coronal | Temporal | 43 | 4.0 | 24.5 | C1 | 23.7 | 25.8 |
|  | T3 | Coronal | Temporal | 159 | 12.2 | 26.8 | C1 | 24.5 | 21.7 |
|  | T4 | Coronal | Temporal | 180 | 12.5 | 28.5 | C1 | 25.1 | 24.4 |

**Table 2.** Comparison of brain strain rates from existing studies. Note that the strain rate in Feng et al. [71] was estimated from Figure 10 in the original article using scheme 1, while the computational scheme for strain rate in Reese et al. [51] was clarified in the caption of Figure 2 in the original article, corresponding to scheme 2 in the current study.

| Scheme | Approach | Study | Loading conditions | Strain rate (s<sup>-1</sup>) |
| --- | --- | --- | --- | --- |
| Scheme 1:<br>Time derivative of strain | Volunteer test | Feng et al. [71] | Linear acceleration: 1.5 g<br>Angular acceleration: 120-140 rad/s<sup>2</sup> | Peak principal strain rate: ~1.2-2.7 |
|  | Animal test | Sullivan et al. [19] | Angular velocity: 121 ± 3.22 rad/s | Mean principal strain rate: 150-189 |
|  | PMHS test | Current study | Linear acceleration: 43-236 g<br>Angular acceleration: 4.0-26.6 krad/s<sup>2</sup><br>Angular velocity: 10-37.7 rad/s | Peak principal strain rate: 9.3-25.8 |
|  | Concussive simulation | Current study | Linear acceleration: 84 g<br>Angular acceleration: 6.2 krad/s<sup>2</sup><br>Angular velocity: 18.8 rad/s | Element-wise maximum principal strain rate: 9.7-143.6<br>Element-wise maximum tract-oriented strain rate: 5.3-114.2 |
| Scheme 2:<br>Rate of deformation tensor | Volunteer test | Reese et al. [51] | Vascular pulsatility | Principal strain rate: 0.1 |
|  | PMHS test | Current study | Linear acceleration: 43-236 g<br>Angular acceleration: 4.0-26.6 krad/s<sup>2</sup><br>Angular velocity: 10-37.7 rad/s | Peak principal strain rate: 15.6-35.6 |
|  | Concussive simulation | Current study | Linear acceleration: 84 g<br>Angular acceleration: 6.2 krad/s<sup>2</sup><br>Angular velocity: 18.8 rad/s | Element-wise maximum principal strain rate: 14.6-119.2<br>Element-wise maximum tract-oriented strain rate: 3.2-63.2 |

**Figure 4.**
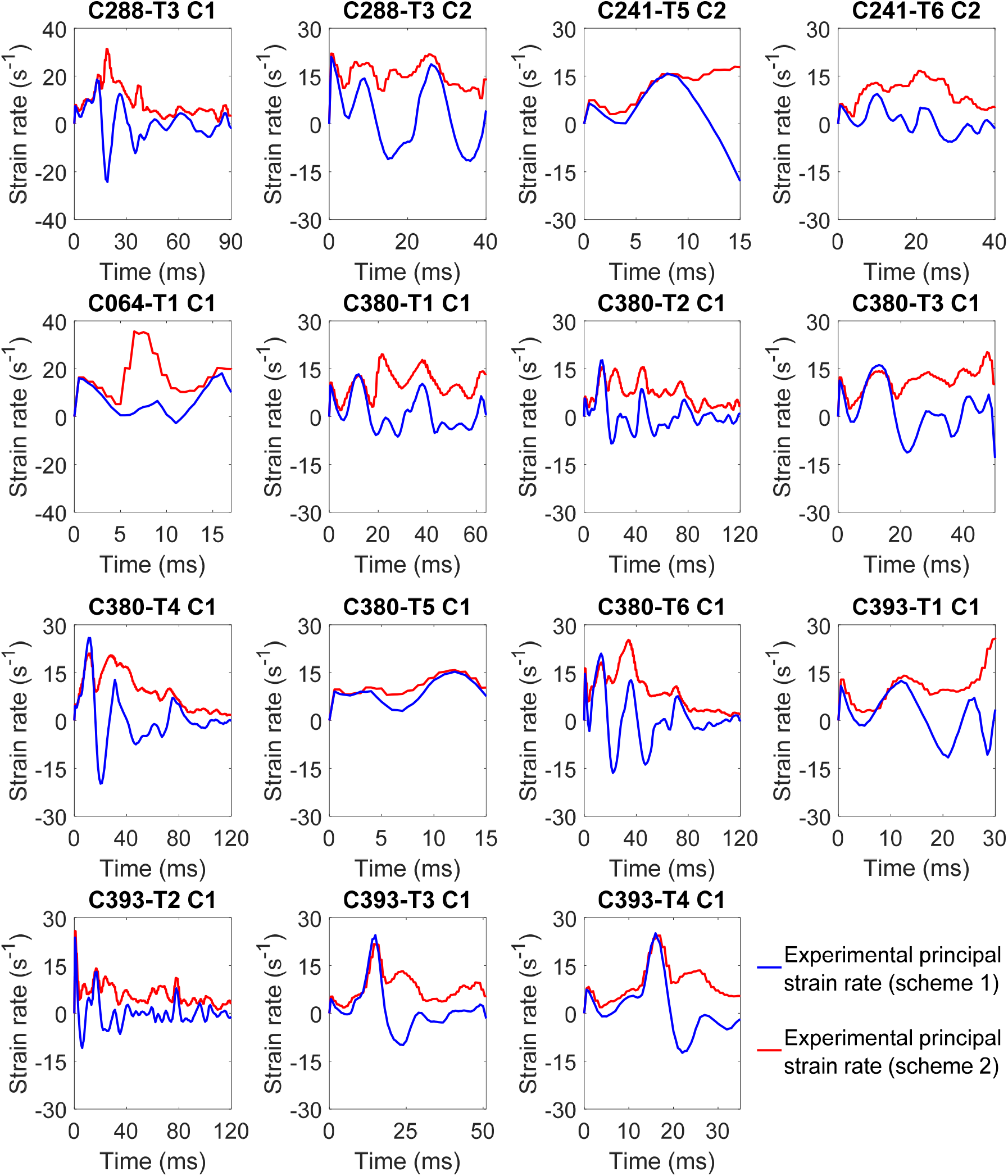
Comparison of experimental principal strain rates computed as time derivate of first principal strain 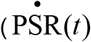 in scheme 1) and first eigenvalue of the rate of deformation tensor 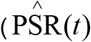 in scheme 2).

### 3.2 Computational principal and tract-oriented strain rates

Figure 5A-F visually compares the maximum principal and tract-oriented strain rates at a representative horizontal cut-section. For the maximum principal strain rate (Figure 5A-C), although both schemes predicted high strain rate peaks (i.e., peaks over or approaching 40 s^-1^) at the cortical region, substantial differences (i.e., differences over 10 s^-1^ or less than -10 s^-1^) between 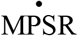 and 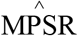 were noted in the left frontal lobe, right occipital lobe, and paraventricular region. For example, 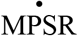 was larger than 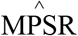 in the left temporal lobe (solid arrow in Figure 5C), while a reverse trend was noted in the left frontal cortex (dash arrow in Figure 5C). When switching to maximum tract-oriented strain rate (Figure 5D-F), 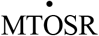 and 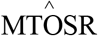 differed in magnitude and spatial distribution. In particular, scheme 1 revealed substantially larger tract-oriented strain rate peaks than scheme 2 across the whole WM in the frontal region (solid arrows in Figure 5F), while 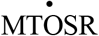 was relatively smaller than 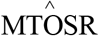 in the external capsules (dash arrows in Figure 5F).

**Figure 5.**
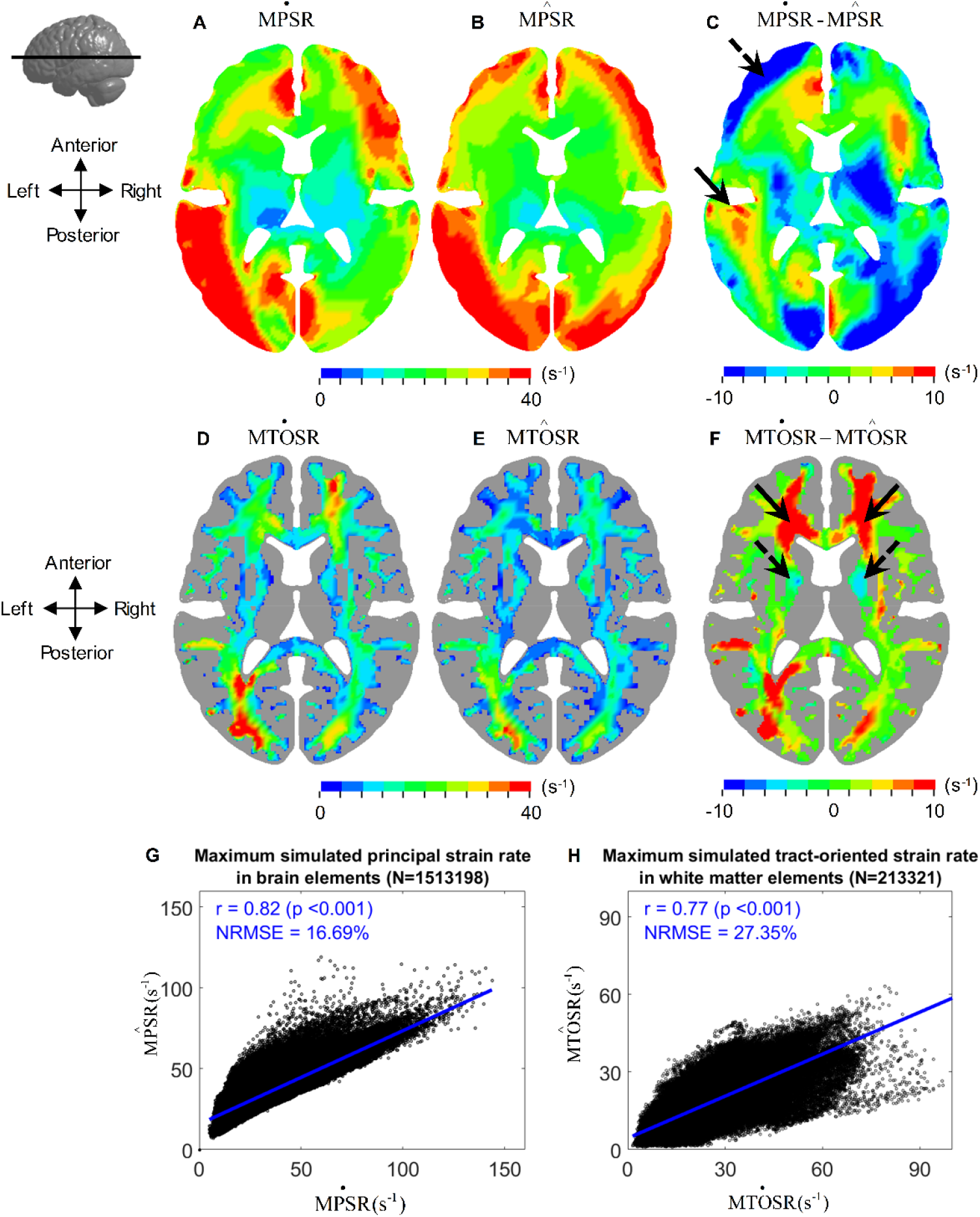
Comparison of time-accumulated maximum principal strain rate and tract-oriented strain rate by both schemes. Upper row: Horizontal cross-sections of 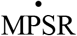 (A), 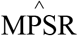 (B), and their relative difference (C). Middle row: Horizontal cross-sections of 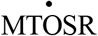 (D), 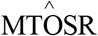 (E), and their relative difference (F). Lower row: Linear regression plots for element-wise maximum principal strain rate in the whole brain (G) and element-wise maximum tract-oriented strain in WM region (H), respectively.

These visible differences were further quantified by linear regression analysis between two schemes (Figure 5G-H). Although a significant correlation (*p*<0.001) was attained for both types of strain rates, the correlation coefficients (r) were 0.82 for maximum principal strain rate and 0.77 for maximum tract-oriented strain rate. Across the whole brain elements (N=1513198), a NRMSE value of 16.69% was computed between 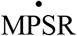 and 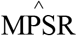, while across the whole WM elements (N=213321), a NRMSE value of 27.35% was calculated between 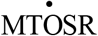 and 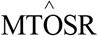.

## 4 Discussion

The current study addressed the ambiguity of brain strain rate computation by presenting a theoretical clarification of two alternative schemes, in which the strain rate was either computed as the time derivative of strain or derived from the rate of deformation tensor. This theoretical disparity was further substantiated by discordant cross-scheme responses in experimental principal strain rate, and computational principal and tract-oriented strain rates. It was thus verified that strain rates were dependent on the specific computational schemes, underscoring that cross-scheme comparison of strain rate results was inappropriate. The newly calculated experimental strain rate curves in the supplementary material, especially those with the peak values attained within the pulse durations, can be used to evaluate the strain rate response of FE models of human head, as exemplified for the ADAPT model in Appendix.

Many experimental models of TBI have reported a strain rate dependency of morphological or functional alterations of experimental tissues secondary to mechanical insults with precisely controlled loading parameters [67-69], while several computational studies have also reported strain rate as an appropriate predictor for brain injury [8, 19, 31, 36, 41]. However, most TBI studies didn’t report details of strain rate calculation, making an appropriate interpretation of strain rate results impossible. For the handful of investigations with the strain rate calculation procedures elaborated, disparate inconsistency was noted in the computational scheme, posing inevitable challenges while comparing strain rate-related findings across studies. Our work demonstrated cross-scheme disparities in the resultant strain rate responses from the theoretical perspective as well as experimental and computational substantiation. These disparities highlighted that cross-scheme comparison of strain rate results is invalid. For example, the strain rate thresholds established by Patton et al. [70] using scheme 2 were not comparable with the strain rate results derived by scheme 1 in the study by Hernandez and Camarillo [45], even though both studies used the exact same FE model [63]. We recommended that, if applicable, clarifying the strain rate computational details be regarded as standard content in future studies.

As both schemes are in use for strain rate computation and no consensus has been reached within the research community, we thus computed the experimental strain rates from the cadaveric tests reported earlier [22] using both schemes. The newly calculated strain rate responses are compared with values reported in other studies (Table 1). When computing the strain rate as the time derivate of strain (i.e., scheme 1), Sullivan et al. [19] estimated the strain rate in six horizontally dissected piglet brains during rapid rotational events, in which the imposed angular velocities were one order of magnitude larger than those in Hardy et al. [22]. Consequently, the reported strain rates in Sullivan et al. [19] largely surpassed the current experimental results. When switching to scheme 2, Reese et al. [51] reconstructed the velocity gradient from phase-contrast magnetic resonance imaging due to the vascular pulsatile and then computed 3D strain-rate tensor of brain parenchyma. This pulsatile-induced strain rate was two orders of magnitude smaller than the current experimental strain rates calculated by scheme 2.

The reliability of FE model-predicted responses can be partially guaranteed only if the computational variables are validated against relevant experimental data. It has been long recognized that an FE head model validated against experimental pressure does not necessarily guarantee an acceptable predictability of the brain displacement or deformation [72-74]. More recently, several independent research groups advocated that a model to be used for strain prediction should be evaluated against the experimental brain strain, and not just the brain-skull relative motion [23, 38, 39]. When it comes to strain rate, there is only one FE model of piglet brain [19], to the best of the authors’ knowledge, having the strain rate peaks (instead of the whole strain rate curves) evaluated. As is the typical practice, FE head models validated against brain-skull relative motion and/or brain deformation are directly used for strain rate prediction. Although the strain rate can be derived from strain or displacement field via a chain of calculation (as detailed in section 2.1), a direct evaluation of FE-derived strain rate responses is preferred if the model is intended to be used for strain rate prediction. This is partially supported by the incongruent strain and strain rate validation performances of the FE model of piglet brain [19], in which the computational strain rate peaks exhibited superior correlation with their experimentally determined counterparts than that between the experimental and computational strain peaks. Note that, in the study by Sullivan et al. [19], the strain rate was computed as the time derivative of strain (i.e., scheme 1 in the current study).

Several recent studies based on FE models of piglet brain [8, 19, 41] have reported that the maximum tract-oriented strain rate exhibited superior injury predictability over maximum principal strain rate, while the majority of the existing FE investigations on human brain remain to solely scrutinize the maximum principal strain rate. We thus examined the relationship between the prevalently used principal strain rate and the recently advocated tract-oriented strain rate across all WM elements in the ADAPT model. As quantitatively illustrated in Figure 6, no consistent scaling from maximum principal strain rate to its tract-aligned counterpart was noted regardless of the computational scheme. This supports the potential incorporation of tract-oriented strain rate as an independent injury metric in future studies of human brain injury.

**Figure 6.**
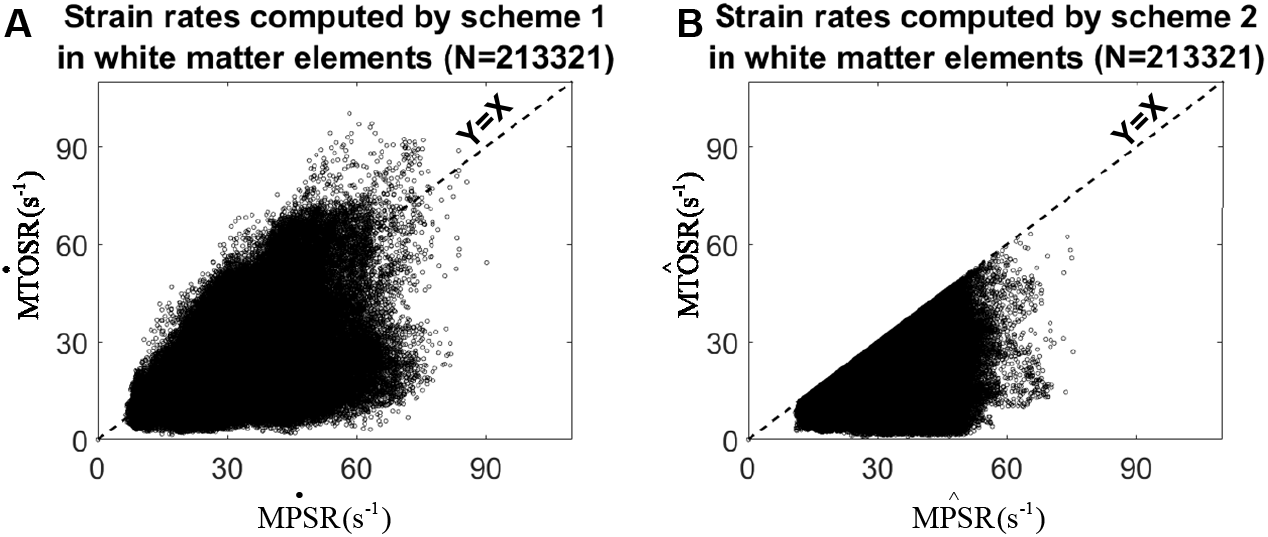
Element-wise comparisons of maximum principal strain rates and tract-related strain rates computed by scheme 1 (A) and scheme 2 (B) based on results of WM elements.

In addition to the strain rate, inconsistency is also noted in strain type in the literature. Even when limiting to TBI-related studies, the adopted strain type includes Green-Lagrange strain [40, 75-78], logarithmic strain (alternatively termed as true strain and natural strain) [79-81], engineering strain [82, 83], incremental strain [22], etc. For a detailed comparison of different strain types, readers are suggested to refer on one study by Karimi and Navidbakhsh [84] or continuum mechanics-related textbooks (e.g., chapter 3 in [85]). It is recommended that the strain type should also be clarified in future studies.

### 4.1 Limitation and future work

Several limitations in this study should be noted, which require further investigation. First, although our results clearly demonstrated cross-scheme disparities, we have no recommendation on which scheme is superior. This can be alternatively done by evaluating the injury predictability of these strain rate-based metrics computed by these two schemes. For the time being, choices may be prioritized differently depending on the intended purpose, but we strongly recommended that the exact strain rate computational schemes be clearly stated. Secondly, the cadaveric experiments analysed in this study were not without errors, as was acknowledged in the original studies [22, 86] and exhaustively evaluated in one recent study [38]. Moreover, the principal strain rate responses estimated from the experimental data and the ADAPT model exhibited certain deviations (Appendix), although this is the first FE human brain model performing strain rate evaluation. Even so, we’d like to highlight that the main advocation of the current work, i.e., cross-scheme comparison of the strain rate results was not valid, is theoretically supported by the content presented in the “Theoretical clarification” section in the current study. Such a advocation should not be disapproved by these experimental and computational substantiations with certain deficits. Nevertheless, we acknowledge that further demonstration of cross-scheme disparity is deemed necessary by employing other experimental data and better validated FE models.

## 5 Conclusion

Given the ambiguity and inconsistency of strain rate computational schemes in existing brain injury studies, the current work presented a theoretical clarification of two rival schemes: strain rates computed as the time derivative of strain and derived from the rate of deformation tensor. Both schemes were employed to calculate experimental principal strain rate and computational principal and tract-oriented strain rates, exhibiting highly scheme-dependent responses. Our work underscores that cross-scheme comparison of strain rate should be avoided and the exact strain rate computational scheme needs to be clarified in future studies. The newly calculated experimental strain rate data in the supplementary material can be used to validate the strain rate response of FE models of human head.

## Acknowledgment

This research has received funding from KTH Royal Institute of Technology (Stockholm, Sweden) and the Swedish Research Council (VR-2020-04496 and VR-2020-04724). The content of this article is solely the responsibility of the authors and does not necessarily represent the official views of funding agencies. The simulations were enabled by resources in projects [NAISS 2023/5-17, SNIC 2022/5-645, SNIC 2022/5-475, SNIC 2021/5-406 and SNIC 2021/5-459] provided by the Swedish National Infrastructure for Computing (SNIC) at the center for High Performance Computing (PDC), partially funded by the Swedish Research Council through grant agreement no. VR-2020-04496 and VR-2020-04724. The authors thank for the two anonymous reviewers and Professor Antoine Jerusalem for the stimulating comments and valuable suggestions that substantially improved this paper. We also acknowledge Dr. Michel Destrade and Dr. Valentina Balbi from University of Galway, Dr. David B MacManus from University College Dublin, Dr. Xiancheng Yu from Imperial College London, Dr. Haojie Mao from Western University, Dr. Rika Wright Carlsen from Robert Morris University, Dr. Declan Patton from Children’s Hospital of Philadelphia, and Dr. T Christian Gasser from KTH Royal Institute of Technology for the helpful discussion on strain rate computation.

## Author contributions

**Zhou Zhou:** Conception and study design, Theoretical investigation, Finite element simulation, Data processing, Visualization; Writing – review & editing; **Xiaogai Li:** Conception and study design, Methodology, Review & editing; **Yuzhe Liu:** Theoretical investigation, Review & editing; **Warren Hardy:** Experimental data resources; **Svein Kleiven:** Conception and study design, Methodology, Review & editing.

## Conflict of Interest

The authors declare that they have no conflict of interest.

## Appendix: Strain rate evaluation of finite element head model

Strain rate responses of the ADAPT model were validated against the newly calculated experimental results. Three representative impacts were selected, including C1 in C288T3 (sagittal impact), C1 in C380T2 (horizontal impact), and C1 in C380T4 (coronal impact). To better reflect the cranial geometry of experimental specimens, the model was scaled to match the reported anthropometric measurements of the cadaveric heads. The nodes nearest to the initial position of the experimental NDT markers were identified and further informed the formulation of the eight tetrahedral elements representing the tissue encased by the experimental NDT cluster. Following the same approach of computing experimental strain rate, the 3D motion of the identified nodes with respect to the skull was acquired from the whole head model simulation and further exercised the tetrahedral elements, through which the first principal strain rate of each element was respectively extracted as the time derivatives of the first principal strain (i.e., 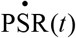 for the scheme 1) and maximum eigenvalue of the rate of deformation tensor (i.e., 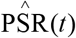 for the scheme 2). For each cluster, the strain rates from all eight elements were averaged.

The correlation between the computationally predicted strain rate responses and experimentally determined measurements was quantitatively evaluated by the cross-correlation method in CORA (correlation and analysis) with technical details available in [87].

Briefly, the cross-correlation method in CORA scored the level of correlation between a pair of time history curves by equally weighting their similarities in shape, size, and phase. The CORA score ranges from 0 (no correlation) to 1 (perfect match). The exact setting of relevant parameters for CORA implementation and biofidelity classification based on CORA scores are respectively presented in Table A1 and Table A2.

**Table A1.** Parameter setting for the cross-correlation in CORA.

| Parameter | Value | Parameter | Value |
| --- | --- | --- | --- |
| A_THRES | 0.030 | INT_MIN | 0.70 |
| B_THRES | 0.075 | K_V | 3 |
| A_VAL | 0.010 | K_P | 1 |
| B_DELTA_END | 0.200 | K_G | 1 |
| T_MIN / T_MAX (ms) | 0/40 | G_V | 0.333 |
| D_MIN | 0.01 | G_P | 0.333 |
| D_MAX | 0.25 | G_G | 0.333 |

**Table A2.** Biofidelity rate and corresponding range of CORA score defined by ISO/TR 9790.

| Biofidelity rating | CORA score |
| --- | --- |
| Excellent | 0.86-1.00 |
| Good | 0.65-0.86 |
| Fair | 0.44-0.65 |
| Margin | 0.26-0.44 |
| Unacceptable | 0-0.26 |

Figure A1 illustrates the comparison of model-predicted strain rate curves against the experiment-derived results for both schemes. The CORA score ranged from 0.65 to 0.71 for scheme 1 and from 0.64 to 0.94 for scheme 2. According to the biofidelity classification based on CORA scores (Table A2), the strain rate response of the ADAPT model can be rated as “good” for scheme 1, and “good” to “excellent” for scheme 2.

**Figure A1.**
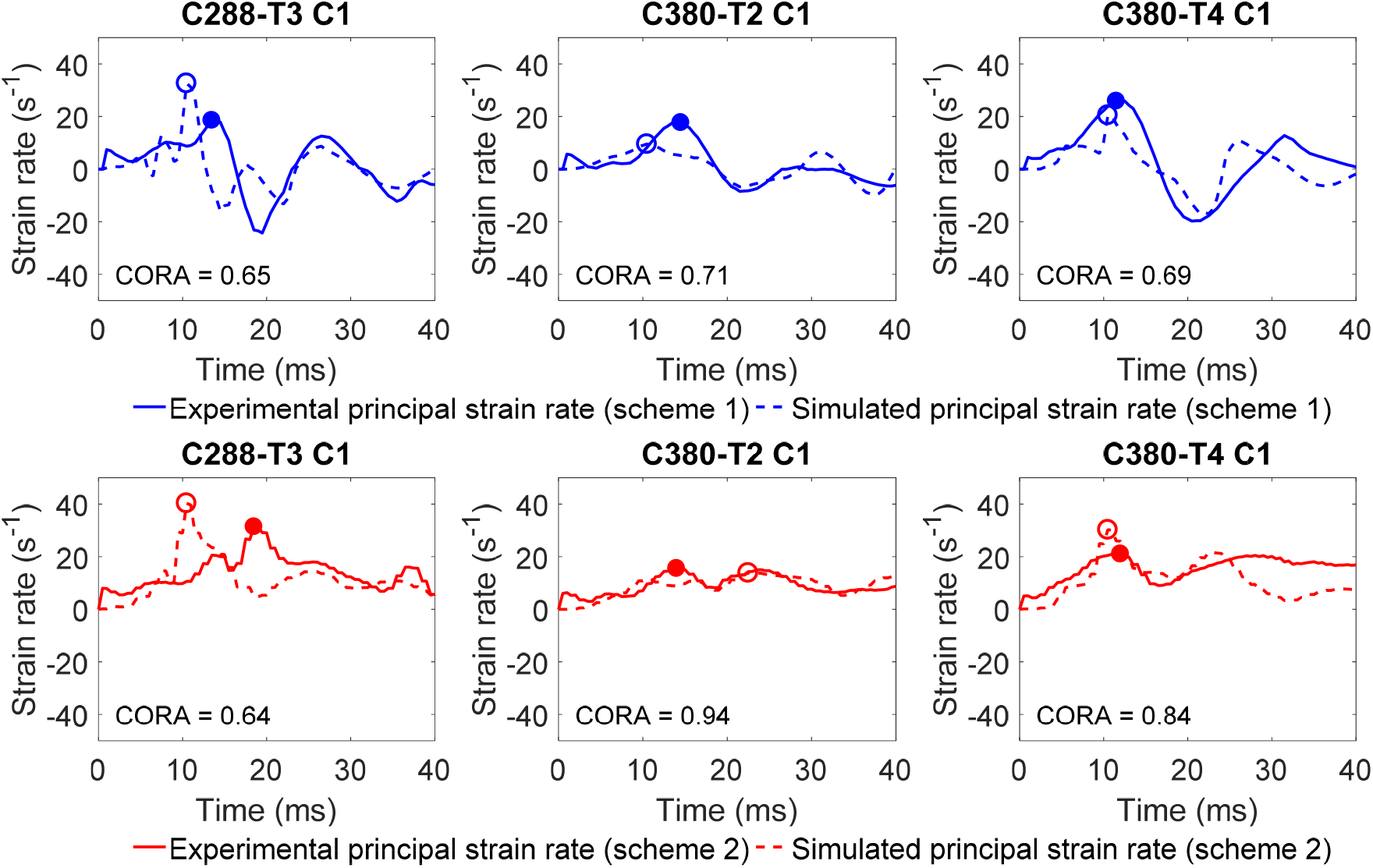
Comparison of experimental and simulated strain rates computed by scheme 1 (upper row) and scheme 2 (low row).

## Notes

### Competing Interest Statement

The authors have declared no competing interest.

### Summary of Updates

see the document

